# Tumor-specific EphA2 receptor tyrosine kinase inhibits anti-tumor immunity by recruiting suppressive myeloid populations in NSCLC

**DOI:** 10.1101/2020.05.08.084830

**Authors:** Eileen Shiuan, Wenqiang Song, Shan Wang, Dana M. Brantley-Sieders, Jin Chen

## Abstract

Given the success of both targeted and immunotherapies against cancer, there is increasing utility for identifying targeted agents that also promote anti-tumor immunity. EphA2 is a receptor tyrosine kinase that contributes to tumor growth and metastasis and has been identified as a viable target for many solid cancers. Investigating EphA2’s impact on the host immune system may advance our understanding of tumor immune evasion and the consequences of targeting EphA2 on the tumor microenvironment. Here, we examine how tumor-specific EphA2 affects the activation and infiltration of immune cell populations and the cytokine and chemokine milieu in non-small cell lung cancer (NSCLC). Effects of EphA2 overexpression in murine NSCLC cells were evaluated in both *in vitro* cell viability assays and *in vivo* tumor models. Tumor immune infiltrate was assessed by flow cytometry. Cytokine and chemokine expression was evaluated using NanoString technology and qRT-PCR. Although EphA2 overexpression in NSCLC cells did not display proliferative advantage *in vitro*, it conferred a growth advantage *in vivo*. Analysis of lung tumor infiltrate revealed decreased natural killer and T cells in the EphA2-overexpressing tumors, as well as increased myeloid populations, including tumor-associated macrophages (TAMs). T cell activation, particularly in CD8 T cells, was decreased, while PD-1 expression was increased. These changes were accompanied by increased monocyte-attracting chemokines, specifically CCL2, CCL7, CCL8, and CCL12, and immunosuppressive proteins TGF-β and arginase 1. Our studies suggest EphA2 on tumor cells recruits monocytes and promotes their differentiation into TAMs that likely inhibit activation and infiltration of cytotoxic lymphocytes, promoting tumor immune escape. Further studies are needed to determine the molecular mechanisms by which EphA2 affects the recruitment of these cell types and to test the function of these myeloid cells.

## Introduction

Targeted and immunotherapies have both emerged as cornerstones of anti-cancer treatment over the past several decades. Despite these advancements, not all patients derive durable clinical benefit from these therapeutic agents, and many efforts are underway to design the best combinations of targeted therapies and immune checkpoint inhibitors (ICIs) that will enhance responses while limiting toxicities^1,2^. However, in order to effectively combine these different treatment modalities, more research is needed to evaluate how one may impact the other – in particular, how targeted agents may already be altering the host anti-tumor immune response.

Despite the introduction of many novel therapies, lung cancer is still the most prevalent cancer that occurs among men and women combined and the leading cause of cancer-related death. A fraction of patients with NSCLC, the most common type of lung cancer, have experienced clinical benefit from single-agent ICIs and/or targeted agents, depending on the mutational profile, but one of the major challenges has been strategizing which combination therapies may be the most synergistic while least toxic and therefore should be tested in trials first^3,4^.

EphA2 receptor tyrosine kinase (RTK) is a cell-surface protein that is overexpressed and implicated in the progression of many solid cancers including NSCLC, and it has become a prominent target for anti-cancer therapy. Currently, agents targeting EphA2, including kinase inhibitors, antibodies, peptide-drug conjugates, liposomal siRNA, chimeric antigen receptor (CAR)-T cells, and dendritic cell (DC) vaccines, are being tested in various cancer clinical trials around the world (NCT02252211, NCT04180371, NCT01591356, NCT01440998, NCT03423992, NCT00563290, NCT00371254, NCT02575261, NCT01876212, NCT00423735). Although the role of EphA2 within the cancer cell and tumor endothelium is well-studied^5,6^, its impact on the tumor immune microenvironment is largely unknown^7^. Thus, we aim to evaluate how EphA2 expression in the tumor affects the immune landscape in NSCLC.

To test the impact of tumor-specific EphA2, we overexpressed EphA2 in murine NSCLC cells and found that EphA2 overexpression increases lung tumor burden *in vivo*. We examined immune infiltrate of tumor-bearing lungs by flow cytometry and found that lungs with EphA2-overexpressing tumors had decreased lymphocytic populations and activation of CD8 T cells but had conversely increased myeloid populations, including TAMs and monocytes, and PD-1 expression in T cells. In addition, gene expression analysis of lung tumors revealed that EphA2 overexpression increased levels of monocyte-attracting chemokines and myeloid-associated immunosuppressive proteins. Together, these studies suggest that EphA2 in NSCLC cells recruits monocytes and promotes their transformation into TAMs, which inhibit the activation of anti-tumor T cells. This provides relevant insight into EphA2’s impact on the immune landscape of NSCLC and the additional potential benefits of targeting EphA2 in cancer.

## Materials and Methods

### Cell culture

Lewis lung carcinoma (LLC) cells were a generous gift from Dr. Barbara Fingleton (Vanderbilt University). *Kras G12D* mutant, *Tp53* and *Stk11/LKB1* knockout, GFP NSCLC cell line (KPL) from the C57BL/6 background was previously generated in the lab^8^. Both cell lines were maintained in DMEM (Corning #MT10013CV) supplemented with penicillin/streptomycin (Gibco #15140163) and 10% FBS (Gibco #A3160502). Luciferase-expressing KPL cells were generated by serial dilutions of KPL cells with lentiviral overexpression of the luciferase gene. Stable EphA2 overexpression was generated by lentiviral transduction of pCDH vector containing the human EphA2 cDNA sequence with subsequent puromycin selection (2 μg/ml) for four days and was compared with empty vector pCDH control.

### Western blotting

Cells were washed with PBS and lysed on ice with RIPA buffer supplemented with a protease inhibitor cocktail (Sigma-Aldrich #4693124001) and phosphatase inhibitors (Sigma-Aldrich #4906845001). Lysates were electrophoresed on a 12% SDS-polyacrylamide gel and transferred to nitrocellulose membranes, which were blocked for a half hour with 5% nonfat dry milk in TBS-T buffer. Membranes were incubated with primary monoclonal antibodies against EphA2 (Cell Signaling Technologies #6997, 1/1000) and tubulin (Sigma-Aldrich #T4026, 1/2000) overnight at 4°C, followed by three washes with TBS-T and incubation with secondary antibodies goat anti-rabbit IRDye 800CW (LI-COR #926-32211) and anti-mouse IRDye 680LT (LI-COR #926-68020, 1/20,000) for 1 hour at room temperature. After washing with TBS-T, blots were imaged using LI-COR Odyssey.

### Cell viability assays

For MTT assays, cells were seeded at a density of 1000 cells per well in 100 μl media in 96-well plates. At each indicated time point, 20 μl of 5 mg/ml thiazolyl blue tetrazolium bromide (MTT) reagent (Thermo Fisher #M6494) in PBS was added per well and incubated at 37°C for 3 hours. Media was then aspirated, and 150 μl of isopropanol solution with 4mM HCl and 0.1% NP-40 was added to each well and rocked at room temperature for 10 minutes. The absorbance at 590 nm was read on a BioTek spectrophotometer and recorded using a Gen5 Microplate Reader. Cell viability was presented as a relative fold change from day 1 values. Each assay included six technical replicates and was repeated at least three independent times. For colony formation assays, cells were seeded at a density of 400 cells per well in 2 ml media in 12-well plates. After incubating for seven days, media was aspirated, and plates were washed x2 with PBS on ice, fixed with methanol for 10 minutes, and stained with 0.5% crystal violet in methanol for 10 minutes at room temperature. Plates were then rinsed under diH2O and let out to dry overnight before image acquisition. Images were analyzed using NIH ImageJ, and colony formation was presented as percentage of total area. Each assay included three technical replicates and was repeated four times.

### Animal models

Animals were housed in a non-barrier animal facility under pathogen-free conditions, 12-hour light/dark cycle, and access to standard rodent diet and water *ad libitum*. Experiments were performed in accordance with AAALAC guidelines and with Vanderbilt University Medical Center Institutional Animal Care and Use Committee approval. Wild-type C57BL/6 mice were purchased from Jackson Laboratory and bred to generate litters for experiments. Male athymic nude (*Foxn1^nu^*) mice were purchased from Envigo for experiments testing tumor growth and immune infiltrate in the context of T cell deficiency. All other experiments utilized male and female immunocompetent wild-type C57BL/6 mice. Mice were co-housed with one to four littermates for at least two weeks prior to and during all experiments and compared with littermate controls whenever possible. All mice used for tumor experiments were six to ten weeks old at the onset on the experiment. Experimental cohorts were limited to litters that were born within two consecutive weeks. Sample sizes are as shown in the figures. At experimental endpoints, mice were euthanized by cervical dislocation.

### Tumor models

For lung tumor colonization experiments, luciferase-expressing KPL cells suspended in PBS were injected via tail vein into seven to ten-week-old wild-type C57BL/6 mice, and mice underwent *in vivo* bioluminescence imaging with a PerkinElmer IVIS Spectrum several hours post-injection to verify successful delivery to the lungs. For experiments comparing lung tumor growth between control and EphA2-overexpressing cells, 1×10^6^ KPL cells were injected in each mouse. For experiments equalizing tumor burden, either 4×10^6^ vector control or 1×10^6^ EphA2-overexpressing KPL cells were injected in each mouse. Mice were reimaged at one and two weeks post-injection and sacrificed at day 14-16. Lungs were weighed, imaged for GFP+ tumors, and processed for flow cytometry analysis. For subcutaneous tumor implantations, 1×10^6^ KPL cells suspended in a 1:1 mixture of PBS and Growth Factor-Reduced Matrigel (Corning #354230) were injected subcutaneously into the dorsal flanks of six to eight-week-old male and female mice. In experiments with athymic nude mice, tumors dimensions were measured by digital caliper at given time points every other day, and volume was calculated using the following formula: volume = length x width^2^ x 0.52. Tumors were subsequently harvested, weighed, and processed for flow cytometry at day 14 post-implantation.

### Flow cytometry

Lungs and subcutaneous tumors were minced and dissociated in RPMI-1640 media (Corning #MT10040CV) containing 2.5% FBS, 1 mg/ml collagenase IA (Sigma-Aldrich #C9891), and 0.25 mg/ml DNase I (Sigma-Aldrich #DN25) for 45 minutes at 37°C. Digested tissue was then filtered through a 70-μm strainer, and red blood cells were lysed using ACK Lysis Buffer (KD Medical #RGF-3015). Samples were washed with PBS and stained with Ghost Dye Violet V450 (Tonbo Biosciences #13-0863) or V510 (Tonbo Biosciences #13-0870) to exclude dead cells. After washing with buffer (0.5% BSA, 2mM EDTA in PBS), samples were blocked in αCD16/32 mouse Fc block (Tonbo Biosciences #70-0161) and stained for extracellular proteins using an antibody master mix made in buffer. After washing with buffer, cells were fixed with 2% PFA. Flow cytometry data was obtained on a BD 4-laser Fortessa using BD FACS Diva software v8.0.1 and analyzed using FlowJo software v10.6.1. Fluorescence minus one (FMO) samples were used as gating controls when needed. Antibodies used in flow panels are detailed in Table 1, and gating strategies used in analysis are detailed in Table 2. Each data point is generated after analyzing at least 5×10^5^ viable cells from a specimen from an individual mouse.

**Table 1.** Antibodies used in flow cytometry analysis.

| Antibody target | Manufacturer | Catalog # | Fluorophore | Dilution | RRID |
| --- | --- | --- | --- | --- | --- |
| MHCII I-E/A | Tonbo Biosciences | 75-5321 | V450 | 1/250 | AB_2621965 |
| CD8a | BD Biosciences | 560469 | V450 | 1/250 | AB_1645281 |
| CD45.2 | Tonbo Biosciences | 35-0454 | V450 | 1/500 | AB_2621965 |
| CD19 | Tonbo Biosciences | 35-0193 | FITC | 1/250 | AB_2621682 |
| CD3e | BD Biosciences | 553061 | FITC | 1/250 | AB_394594 |
| CD44 | Tonbo Biosciences | 35-0441 | FITC | 1/250 | AB_2621688 |
| NK1.1 | BD Biosciences | 557391 | PE | 1/250 | AB_396674 |
| CD11b | eBiosciences | 12-0112-82 | PE | 1/500 | AB_2734869 |
| CD25 | eBiosciences | 12-0251-81 | PE | 1/500 | AB_465606 |
| CTLA-4 | BD Biosciences | 561718 | PE | 1/250 | AB_10895585 |
| CD3e | BD Biosciences | 561826 | APC | 1/500 | AB_10896663 |
| CD4 | Biologend | 100516 | APC | 1/1000 | AB_312719 |
| F4/80 | eBiosciences | 17-4801-82 | APC | 1/250 | AB_2784648 |
| CD3e | Tonbo Biosciences | 65-0031 | PerCP-Cy5.5 | 1/250 | AB_2621872 |
| CD11b | Tonbo Biosciences | 65-0112 | PerCP-Cy5.5 | 1/500 | AB_2621885 |
| CD11c | BD Biosciences | 560584 | PerCP-Cy5.5 | 1/250 | AB_1727422 |
| F4/80 | eBiosciences | 45-4801-82 | PerCP-Cy5.5 | 1/250 | AB_914345 |
| Ly6C | eBiosciences | 45-5932-82 | PerCP-Cy5.5 | 1/250 | AB_2723343 |
| Ly6G(Gr1) | eBiosciences | 45-5931-80 | PerCP-Cy5.5 | 1/500 | AB_906247 |
| Ly6G(Gr1) | Tonbo Biosciences | 80-5931 | rF710 | 1/1000 | AB_2621999 |
| CD8a | Tonbo Biosciences | 80-0081 | rF710 | 1/500 | AB_2621977 |
| MHCII I-E/A | Biologend | 107622 | Ax700 | 1/1000 | AB_493727 |
| PD-1 | BD Biosciences | 565815 | APC-R700 | 1/500 | AB_2739366 |
| CD4 | Tonbo Biosciences | 55-0041 | PE-Cy5 | 1/2500 | AB_2621816 |
| CD11b | Tonbo Biosciences | 55-0112 | PE-Cy5 | 1/5000 | AB_2621818 |
| CD3e | BD Biosciences | 552774 | PE-Cy7 | 1/750 | AB_394460 |
| NKp46 | eBiosciences | 25-3351-82 | PE-Cy7 | 1/750 | AB_2573442 |
| CD11c | BD Biosciences | 561022 | PE-Cy7 | 1/500 | AB_2033997 |
| CD69 | BD Biosciences | 552879 | PE-Cy7 | 1/500 | AB_394508 |
| CD25 | Tonbo Biosciences | 60-0251 | PE-Cy7 | 1/500 | AB_2621843 |
| CD45 | BD Biosciences | 557659 | APC-Cy7 | 1/500 | AB_396774 |

**Table 2.** Gating strategy used in flow cytometry analysis.

| Cell population | Gating strategy |
| --- | --- |
| KPL cells | CD45-, GFP+, SSA<sup>hi</sup> |
| Immune cells | CD45+, GFP- |
| NK cells (nude mice) | CD45+, GFP-, CD19-, NKp46+ |
| B cells (nude mice) | CD45+, GFP-, CD19+, MHCII+ |
| Gr1+ myeloid cells (nude mice) | CD45+, GFP-, CD11b<sup>hi</sup>, Ly6G(Gr1)+, MHCII- |
| Gr1- myeloid cells (nude mice) | CD45+, GFP-, CD11b<sup>hi</sup>, Ly6G(Gr1)- |
| Macrophages (nude mice) | CD45+, GFP-, CD11b<sup>hi</sup>, F4/80+, Ly6G(Gr1)- |
| DCs (nude mice) | CD45+, GFP-, CD11c+, MHCII+, NKp46-, Ly6G(Gr1)- |
| CD4 T cells | CD45+, GFP-, CD3e+, NK1.1-, CD4+, CD8a- |
| CD8 T cells | CD45+, GFP-, CD3e+, NK1.1-, CD4-, CD8a+ |
| NK cells | CD45+, GFP-, CD3e-, NK1.1+ |
| Macrophages | CD45+, GFP-, CD11b<sup>hi</sup>, F4/80+, Ly6C- |
| Monocytes | CD45+, GFP-, CD11b<sup>hi</sup>, F4/80-, Ly6C+ |
| DCs | CD45+, GFP-, CD11c+, MHCII+, F4/80- |

### NanoString nCounter assay

Lung tumors were dissected and minced with surgical tools cleaned with RNaseZAP (Sigma-Aldrich #R2020), and RNA was extracted using RNeasy Micro Kit (Qiagen #74004). RNA concentration was assessed with a Nanodrop spectrophotometer, and RNA quality was determined by the Agilent 2100 Bioanalyzer System. 20 ng of RNA from each of twelve mice, six gender-matched littermate pairs, were used for input into nanoString nCounter hybridization and hybridized to the nanoString nCounter Mouse PanCancer Immune Profiling Panel probeset (nanoString Technologies #XT-CSO-MIP1-12) to measure gene expression. Raw count data was normalized by background correction, positive control correction, and housekeeping gene correction and log2 transformed using nanoString’s nSolver software v3.0. This software was also used to generate pathway scores, differential gene expression analysis, and the volcano plot. Heatmap was generated with normalized data standardized by gene using Microsoft Excel 2016.

### RT-PCR

RNA was isolated and measured as detailed above and then converted to cDNA using iScript cDNA Synthesis Kit (Bio-Rad #1708891). Quantitative RT-PCR was carried out with TaqMan Gene Expression Assay reagents, specifically TaqMan Fast Advanced Master Mix (Thermo Fisher #4444557) and probes (Table 3), using a StepOnePlus RT-PCR system (Applied Biosystems). Reactions were run in triplicate and mouse *Actb* was used as a housekeeping gene. Six gender-matched littermate pairs were used to validate nanoString hits, and data was presented as fold change of EphA2-overexpressing tumor samples with respect to their littermate control sample.

**Table 3.** TaqMan RT-PCR probes.

| Target gene | Assay ID |
| --- | --- |
| *Actb* | Mm02619580_g1 |
| *Arg1* | Mm00475988_m1 |
| *C5ar1* | Mm00500292_s1 |
| *Ccl2* | Mm00441242_m1 |
| *Ccl7* | Mm00443113_m1 |
| *Ccl8* | Mm01297183_m1 |
| *Ccl12* | Mm01617100_m1 |
| *Ccr2* | Mm99999051_gH |
| *Csf1r* | Mm01266652_m1 |
| *Tgfb1* | Mm01178820_m1 |
| *Tgfb2* | Mm00436955_m1 |
| *Tgfb3* | Mm00436960_m1 |

### Statistical analysis

All graphs and statistical analysis were completed using GraphPad Prism software v6.07. For comparisons of continuous variables between two groups, an unpaired, two-tailed student’s t test with Welch correction or unpaired Mann-Whitney *U*-test was performed. For comparisons of continuous variables over time between two groups, a two-way analysis of variance (ANOVA) was performed. For comparison of survival curves, a log-rank test was performed. For RT-PCR analysis, a one-sample Wilcoxon signed rank test was performed. A *P*-value less than 0.05 was considered statistically significant.

## Results

### EphA2 confers growth advantage to NSCLC in vivo but not in vitro

Because EphA2 has been shown to play both tumor-promoting and suppressing roles, we investigated whether EphA2 overexpression in our murine NSCLC cell lines could impact cell viability and tumor growth. We chose two NSCLC cell lines originating from the C57BL/6 background, LLC and a KRAS G12D mutant cell line containing loss of TP53 and STK11/LKB1 (KPL)^8^, both of which are common alterations found concomitantly with KRAS mutations in human NSCLC samples^9,10^. We stably overexpressed EphA2 in these cells (Figure 1A) but found no changes in cell viability by MTT or colony formation assays *in vitro*, compared to vector control cells (Figure 1B, 1C). We next determined if EphA2 overexpression could impact tumor growth *in vivo* using two different models, subcutaneous implantation and tail vein injection for generation of lung tumors in immunocompetent, wild-type C57BL/6 mice. In both subcutaneous and lung tumor models, EphA2-overexpressing KPL cells grew dramatically larger tumors *in vivo*, as shown by bioluminescence imaging and gross lung specimens (Figure 1D-F). In addition, survival of tail vein-injected mice was significantly worse in mice injected with EphA2-overexpressing cells, compared to control cells (Figure 1G). Overall, these data suggest host factors contribute to EphA2’s impact on *in vivo* growth, which is not apparent *in vitro*.

**Figure 1.**
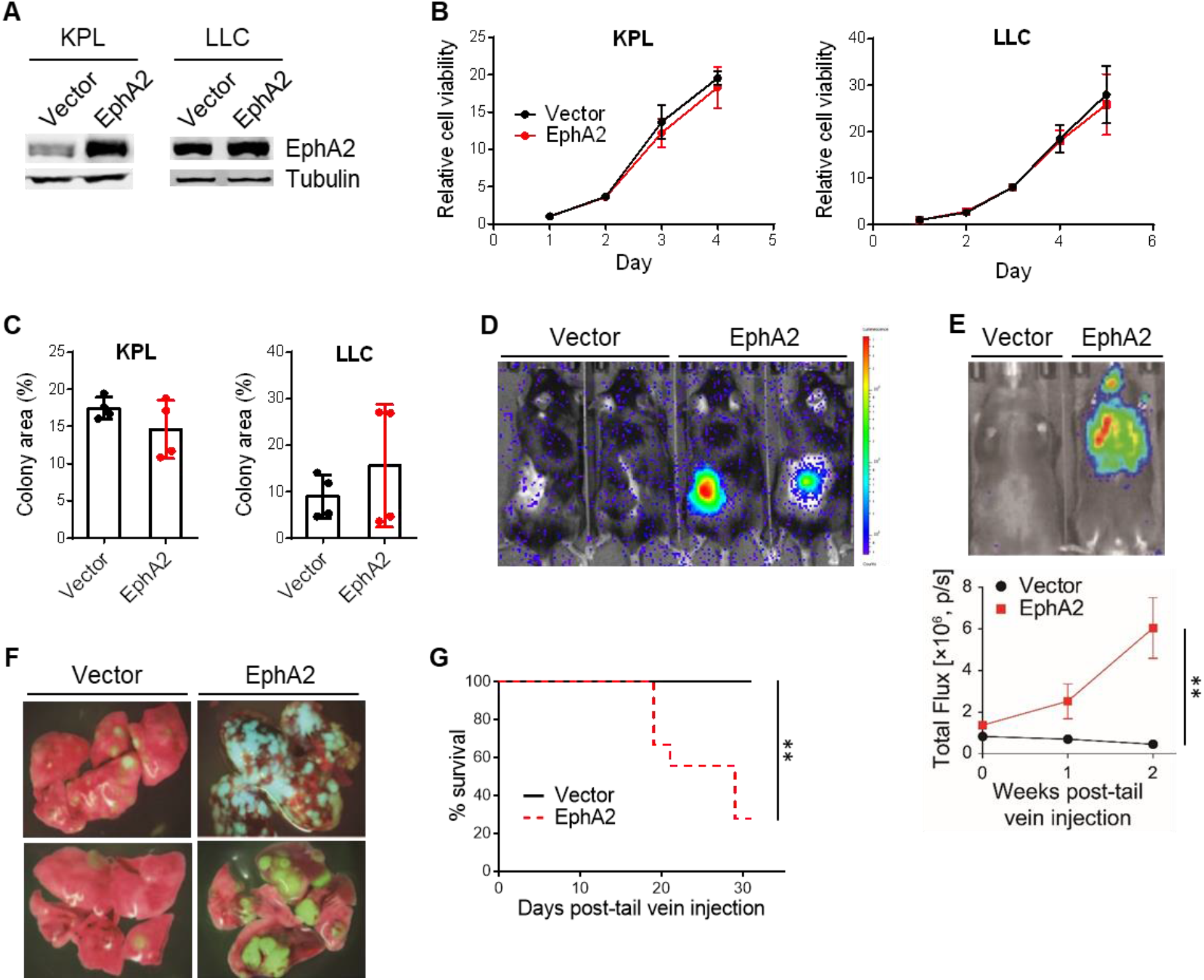
EphA2 confers growth advantage to NSCLC *in vivo* but not *in vitro*. (A) Confirmation of EphA2 overexpression in KPL and LLC cells by western blot. (B, C) *In vitro* cell viability of KPL and LLC cells with control and EphA2 overexpression by MTT and colony formation assays (n=4). (D) Representative image of bioluminescence signal in control and EphA2-overexpressing KPL tumors 14 days after subcutaneous implantation. (E) Representative image of bioluminescence signal 14 days after tail vein injection of control and EphA2-overexpressing KPL cells and quantification of bioluminescence signal at indicated time points (\*\**p*<0.01, two-way ANOVA) (F) Representative gross specimens of GFP+ vector and EphA2-overexpressing KPL tumor-bearing lungs. (G) Survival of mice injected with vector or EphA2-overexpressiong KPL cells via tail vein (\*\**p*<0.01, log-rank test). Data shown are averages ± SD.

### EphA2 overexpression in NSCLC does not significantly impact tumor burden or immune infiltration in nude mice

There are various host factors that may explain the discrepancy between our *in vitro* and *in vivo* results regarding the impact of EphA2 in the NSCLC cell, one of which includes the host immune system. Besides one study demonstrating that tumor-intrinsic EphA2 can inhibit anti-tumor immunity in pancreatic cancer^11^, the effect of EphA2 on the tumor immune microenvironment has been largely unstudied. Thus, we set out to determine if the host immune system plays a role in EphA2-mediated KPL tumor growth *in vivo*. To test whether the adaptive or innate immune response may play a role, we first evaluated KPL tumor growth in athymic nude mice, which have defective T cell adaptive immunity. We did not observe significant differences in tumor growth and weight between control and EphA2-overexpressing tumors in nude mice (Figure 2A). In addition, we performed flow cytometry analysis on the tumors and draining lymph nodes and found no differences in percentages of GFP+ KPL tumor cells and tumor-infiltrating immune cells (Figure 2B). We also observed no significant differences in subsets of immune cells, including natural killer (NK) cells, B cells, DCs, and myeloid cells, from both the tumors and draining lymph nodes in the nude mice (Figure 2C, 2D). These results suggest that EphA2-mediated KPL tumor growth *in vivo* may require suppression of T cell adaptive immunity.

**Figure 2.**
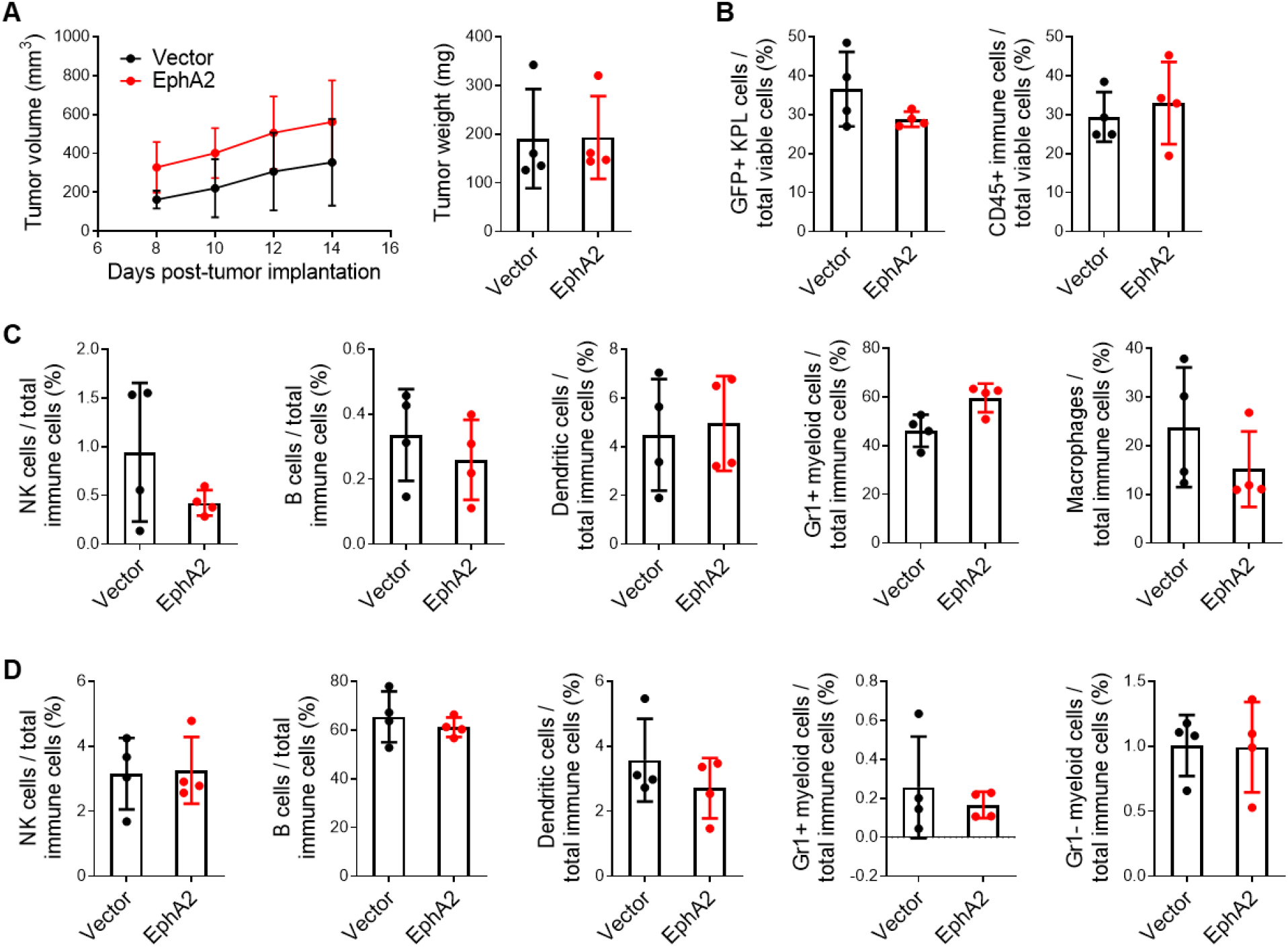
EphA2 overexpression in NSCLC does not significantly impact tumor burden or immune infiltration in nude mice. (A) Tumor volumes over time and weights on day 14 post-implantation of control and EphA2-overexpressing KPL subcutaneous tumors from nude mice. (B) Flow cytometric analysis of GFP+ KPL tumor cells and total tumor-infiltrating immune cells, as well as (C) tumor-infiltrating NK cells, B cells, DCs, macrophages, and Gr1+ myeloid cells. (D) Similar flow cytometry analysis of immune populations from draining inguinal lymph nodes. Data shown are averages ± SD (n=4 mice per group).

### EphA2 overexpression in NSCLC decreases lymphocytic and increases myeloid infiltrate in tumor-bearing lungs

We returned to our tumor model with immunocompetent C57BL/6 mice to examine the tumor immune infiltrate, including the T cell populations. As before, wild-type mice were injected via tail vein with control or EphA2-overexpressing KPL cells, and lungs were harvested for flow cytometry analysis 14 days later. EphA2-overexpressing KPL tumor-bearing lungs had significantly higher percentages of GFP+ KPL tumor cells compared to control (Figure 3A). Although the overall percentage of immune cells did not differ (Figure 3A), the lymphocytic populations, particularly CD4 and CD8 T cells and NK cells, were significantly decreased in the EphA2-overexpressing tumor-bearing lungs (Figure 3B). Conversely, myeloid populations, including macrophages and monocytes, were increased, while there was no change in DCs (Figure 3C). This data suggests that EphA2 in the tumor cell reshapes the tumor immune microenvironment of the lung and shifts it from a lymphocytic to a more myeloid response. This myeloid response presumably plays a greater tumor-promoting role and may suppress the anti-tumor response primarily led by the T cells.

**Figure 3.**
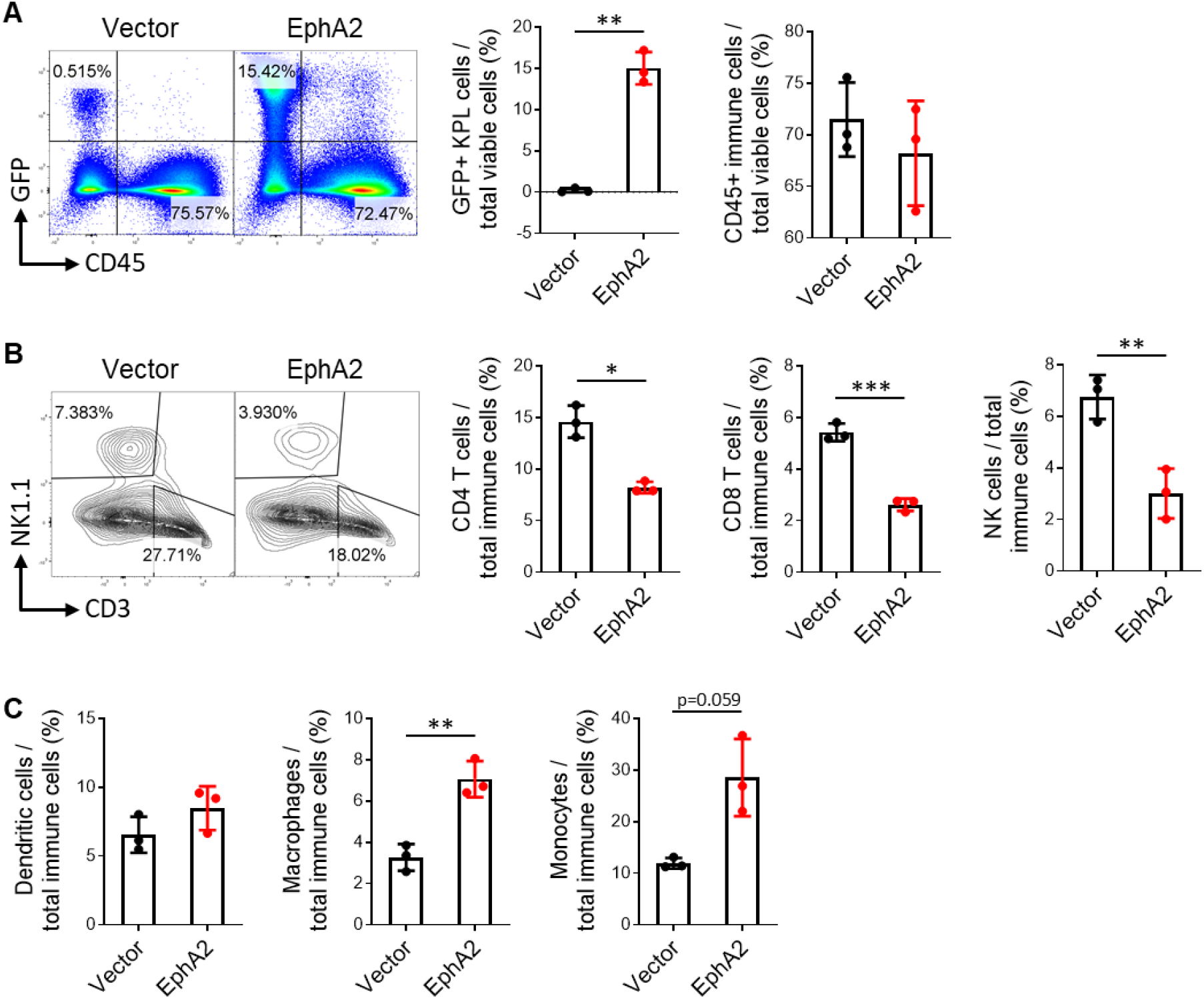
EphA2 overexpression in NSCLC decreases lymphocytic and increases myeloid infiltrate in tumor-bearing lungs. (A) Representative flow cytometry plots and quantification of GFP+ KPL and immune cells from vector control and EphA2-overexpressing tumor-bearing lungs on day 14 post-tail vein injection. (B) Similar flow plots and analysis of CD4 and CD8 T cells and NK cells, as well as (C) quantification of DCs, macrophages, and monocytes. Data shown are averages ± SD (n=3 mice per group, \**p*<0.05; \*\**p*<0.01; \*\*\**p*<0.001, two-tailed unpaired student’s t test with Welch correction).

### EphA2 overexpression in NSCLC suppresses tumor-infiltrating T cells

Although we found a decrease in lymphocyte populations in EphA2-overexpressing tumor-bearing lungs, this result could also be explained by the dramatically greater tumor burden in these lungs, compared to the control lungs, which contained less than a percentage of KPL tumor cells (Figure 3A). Many previous studies have shown a correlation between tumor bulk and the quality of immune infiltrate, though it is difficult to discern which is the cause and which is the effect. In order to determine if tumor bulk contributes to the immune landscape we observe in Figure 3, we repeated the tail vein experiment controlling for tumor bulk by injecting four times the number of control KPL cells, compared to EphA2-overexpressing KPL cells. By day 16, there was comparable amount of KPL tumor burden in control and EphA2-overexpressing tumor-bearing lungs (Figure 4A). Flow cytometric analysis on these lungs did not recapitulate results from the previous experiment, specifically no decrease in lymphocyte populations in EphA2-overexpressing tumor-bearing lungs (Figure 4B). Interestingly, there was an increase in CD4 T cells in the EphA2-overexpressing tumor-bearing lungs. These findings indicate that the differences in immune lung infiltrate we previously observed were at least partially due to increased KPL tumor burden from EphA2-overexpressing tumors compared to control tumors.

**Figure 4.**
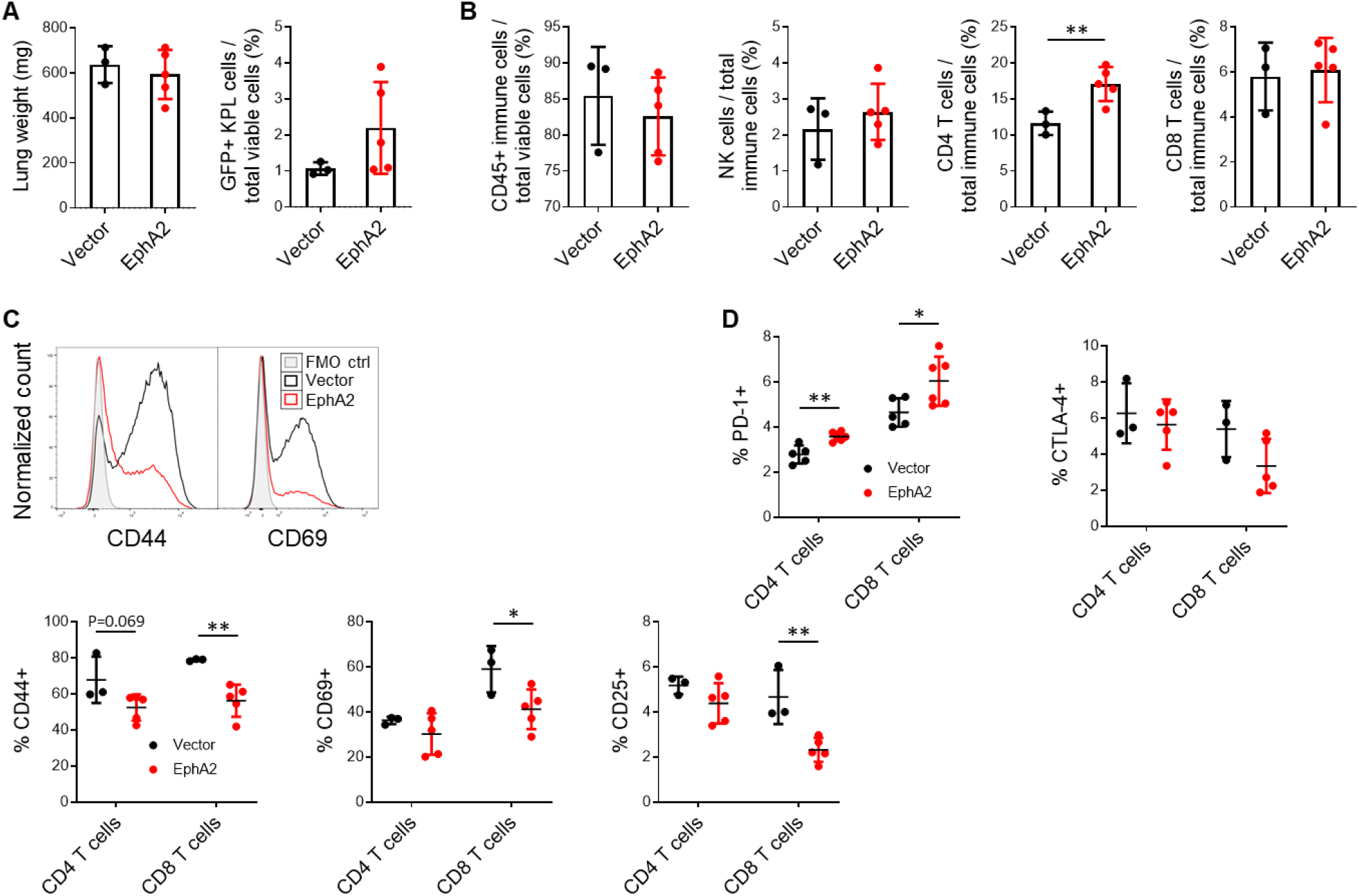
EphA2 overexpression in NSCLC suppresses tumor-infiltrating T cells. (A) Lung weights and quantification of GFP+ KPL cells via flow cytometry from vector control and EphA2-overexpressing tumor-bearing lungs with equalized tumor burden. (B) Flow cytometric analysis of total immune cells, CD4 and CD8 T cells, and NK cells in KPL-tumor bearing lungs. (C) Representative flow histograms of CD44 and CD69 expression on CD8 T cells and quantification of CD44, CD69, and CD25 activation markers on CD4 and CD8 T cells. (D) Quantification of PD-1 and CTLA-4 exhaustion markers on CD4 and CD8 T cells. Data shown are averages ± SD (n=3-6 mice per group, \**p*<0.05; \*\**p*<0.01, two-tailed unpaired student’s t test with Welch correction).

Although the proportion of CD8 T cells did not differ when tumor burden between control and EphA2-overexpressing KPL tumors were equalized, T cell activation status and effector function may still be different. Tumor-infiltrating T cells with upregulated expression of activation markers, such as CD44, CD69, and CD25, and downregulated expression of exhaustion markers like PD-1 and CTLA-4 indicate a higher T cell functional status that mediates a stronger and more enduring anti-tumor response^12^,^13^. We examined the expression of activation markers of T cells from KPL tumor-bearing lungs and found that EphA2 overexpression in the tumor significantly downregulates surface activation markers CD44, CD69, and CD25 on CD8 T cells (Figure 4C). Similar trends were also seen in CD4 T cells, though not significant (Figure 4C). In addition, tumor-specific EphA2 overexpression led to increased PD-1 expression in both CD4 and CD8 T cells, though not CTLA-4 (Figure 4D). These results indicate that EphA2 overexpression in KPL tumors inhibits CD8 T cell activation and may promote T cell exhaustion in the lung tumor microenvironment.

### Gene expression profiling reveals higher expression of myeloid markers and chemoattractants in EphA2-overexpressing tumors

In order to understand the mechanism by which tumor-specific EphA2 suppresses CD8 T cell activation, we performed gene expression profiling of dissected control and EphA2-overexpressing lung tumors using nanoString’s Mouse PanCancer Immune Profiling Panel. Gene pathway analysis yielded two significantly upregulated pathways in the EphA2-overexpressing tumors, cancer progression and macrophage functions, consistent with our observations on *in vivo* tumor burden and findings from flow cytometry (Figure 5A, 5B). Differential expression analysis of individual genes revealed 95 genes that were significantly different in expression between vector control and EphA2-overexpressing tumors with a log2ratio less than −0.5 or greater than 0.5 (Figure 5C).

Among these genes included a group of monocyte and macrophage surface markers (*Csf1r, Ccr2, Itgam*/CD11b, *Mrc1/CD206*), myeloid chemoattractants (*Ccl2, Ccl7, Ccl8, Ccl12*), and immunosuppressive proteins (*Arg1, Tgfb1, Tgfb2, Tgfb3*) (Figure 5D). CD11b is a nonspecific myeloid marker, while CCR2, CSF1R, and CD206 are typically expressed in inflammatory monocytes and macrophages, with CD206 specifically detected in tumor-promoting M2 macrophages^14,15^. CCL2, CCL7, CCL8, and CCL12 all belong to the CCL chemokine family and have been shown to bind to CCR2 expressed on circulating monocytes and facilitate their recruitment^14–16^. Additionally, arginase 1 (ARG1) and TGF-β are well-known immunosuppressive proteins that are secreted in the tumor microenvironment by tumor-promoting cells such as M2 TAMs and subsequently inhibit CD8 T cell effector function^14,15^. We validated a majority of these genes using RT-PCR (Figure 5E).

**Figure 5.**
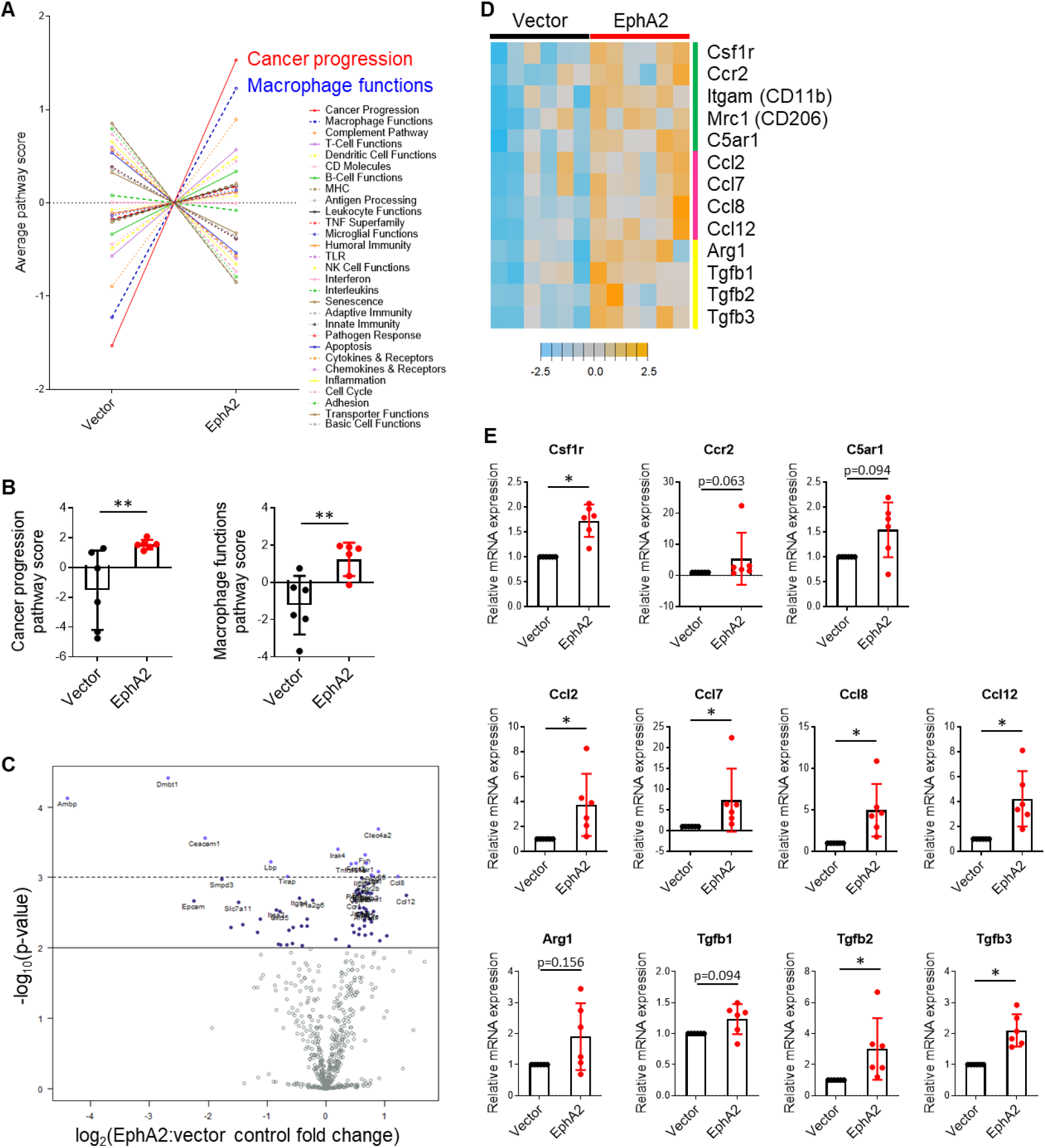
Gene expression profiling reveals higher expression of myeloid markers and chemoattractants in EphA2-overexpressing tumors. (A) Average pathway scores of vector control and EphA2-overexpressing KPL tumors calculated from normalized gene expression data using nanoString nSolver software. (B) Comparison of cancer progression and macrophage functions pathway scores between control and EphA2-overexpressing samples. (n=6 mice per group, \*\**p*<0.01, unpaired Mann-Whitney test) (C) Volcano plot of statistically significant differentially expressed genes. (D) Heatmap depicting standardized expression of differentially expressed myeloid markers (green bar), myeloid-attracting chemokines (pink bar), and immunosuppressive proteins (yellow bar). (E) RT-PCR validation of nanoString hits. (n=6 mice per group, \**p*<0.05, one-sample Wilcoxon signed rank test). Data shown are averages ± SD.

In summary, gene expression analysis provided evidence that tumor-specific EphA2 increases the levels of myeloid chemoattractants, monocyte/macrophage lineage cells, and immunosuppressive proteins. We propose that EphA2 in the tumor cell upregulates expression of chemokines in the tumor milieu that recruit circulating monocytes into the tumor, where they differentiate into macrophages and are co-opted to serve tumor-promoting functions as polarized M2 TAMs (Figure 6). These tumor-promoting functions include secretion of proteins, such as arginase and TGF-β, that inhibit the expansion and activity of T cells, particularly anti-tumor CD8 cytotoxic lymphocytes. This leads to a shift towards a more myeloid and less lymphocytic infiltrate, as we observed in our flow cytometry studies, as well as decreased activation and increased exhaustion in T cells. Ultimately, the effect of EphA2 in the tumor dampens the anti-tumor immune response and perpetuates the vicious cycle of tumor immune escape and growth.

**Figure 6.**
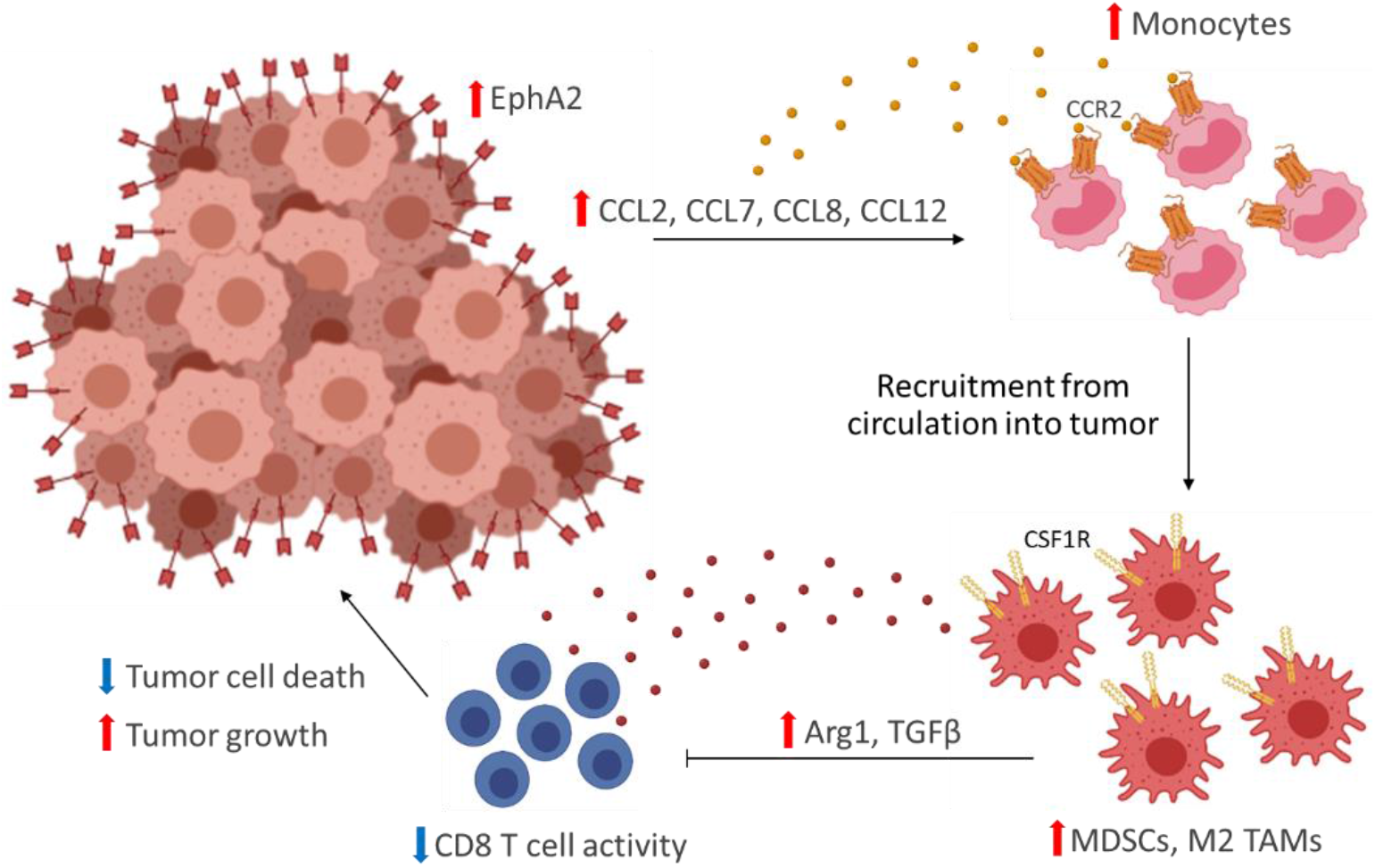
Proposed role of tumor-specific EphA2 on promoting NSCLC immune evasion.

## Discussion

Our studies identify a novel mechanism that contributes to EphA2’s pro-tumor effect in NSCLC. Although EphA2 has previously been shown to facilitate lung tumor growth and metastasis through tumor cell-intrinsic mechanisms^17,18^, this is the first study that we are aware of that shows an immune-mediated phenotype. Despite these new insights, this work poses several unanswered questions. First, which cell types are critical for the EphA2-mediated effect on CD8 T cells? EphA2 overexpression in the cancer cell may directly upregulate myeloid-attracting chemokine expression in the tumor cells, but it may also indirectly affect expression in other stromal cells that can secrete these chemoattractants. Similarly, although we propose that the monocytes and macrophages in EphA2-overexpressing tumors are responsible for the higher levels of arginase and TGF-β, these immunosuppressive proteins can also be secreted from cancer or stromal cells. Further studies utilizing single-cell techniques will be required to elucidate the roles of each cell type in the tumor microenvironment. Additionally, the molecular mechanisms by which tumor-specific EphA2 alters the chemokine milieu and immune landscape of lung cancer may be very complex. EphA2 in the tumor cell can signal in an ephrin ligand-dependent or independent manner^19^, and it can even be packaged into extracellular vesicles to initiate signaling from a distance^20^. Further molecular investigations are needed to better understand the signaling modalities and downstream pathways in the cancer cell that mediate this immune modulation.

Our findings are consistent with a recently published paper that demonstrated tumor cell-intrinsic EphA2 inhibits anti-tumor immunity in pancreatic cancer through regulation of *PTGS2*, the gene encoding cyclooxygenase-2 (COX2)^11^. Markosyan, et al. identified EphA2 from a screen of The Cancer Genome Atlas (TCGA) dataset evaluating genes that were inversely correlated with *CD8A* expression in human pancreatic adenocarcinomas, as well as other markers of cytotoxic T lymphocytes (CTLs). They went on to evaluate the effect of knocking out tumor cell-specific EphA2 in a mouse model of pancreatic cancer bearing a *Kras G12D* mutation, loss of *Tp53*, and a YFP marker (KPCY) and found that EphA2 knockout significantly increased T cell infiltration and activation, as we had observed in our murine KPL model of NSCLC. Interestingly, they found that EphA2 knockout decreased numbers of granulocytic MDSCs but did not affect macrophages. In addition, these authors identified *Ptgs2* from a RNAseq experiment examining the differentially expressed genes between EphA2 WT and KO tumors.

We can draw many parallels between our investigations and the studies performed by Markosyan et al.; however, there are also some intriguing differences. Although both studies demonstrate that tumor cell-specific EphA2 has a detrimental impact on T cell-mediated immunity, one suggests that granulocytic myeloid cells play an intermediary role, while the other suggest monocytic myeloid cells. In addition, *Ptgs2* was not among the genes we found to be differentially expressed between control and EphA2-overexpressing KPL tumors. Furthermore, Markosyan et al. observed a decrease in tumor cell proliferation and *in vivo* tumor burden with EphA2 KO, but we observed no differences in growth *in vitro* or *in vivo* when we knocked out EphA2 in our KPL model via CRISPR/Cas9 (data not shown). Lastly, although the authors observed inverse correlations between *EPHA2* and CTL gene signatures in human pancreatic cancer, we did not discover any consistent trends between *EPHA2* and *CD3E, CD8B, GZMB, PRF1*, or *IFNG* expression from the TCGA lung adenocarcinoma dataset (data not shown). These discrepancies could be due to differences in cancer cell type and model and the EphA2 receptor/ephrin ligand balance in the tumor microenvironment, which can be very dissimilar between the pancreas and lung. Although EphA2 is implicated in tumor growth and metastasis of many types of solid cancers through ligand-independent signaling^21^,^22^, it can also suppress cell growth via ligand-dependent signaling through ephrin-A1, its primary binding partner^23–25^. Because we observed no effect on cell viability *in vitro* or tumor growth *in vivo* with EphA2 knockout in our KPL lung model, perhaps EphA2 is playing a dual tumor promoter and suppressor role in this cell line. Whereas, in KPCY pancreatic tumors, EphA2 is more of a tumor promoter than suppressor. This would suggest that EphA2 is signaling differently in these two tumor types, or at least in these two particular models, and may explain why the immune phenotypes differ.

A note of caution that we feel compelled to point out is that our investigations rely heavily on an overexpression system, which has been shown in published studies to yield artifactual results^26–31^ and likely also observed in many unpublished works. We used the same lentiviral vector to overexpress other genes in KPL cells, including a catalytically dead Cas9 (dCas9), and found that regardless of the gene, overexpression appeared to increase tumor burden *in vivo*, compared to vector control (data not shown). We then tried to overexpress EphA2 using a retroviral vector and discovered that this unfortunately did not recapitulate results we observed in tumor burden with the lentiviral vector (data not shown), though we did not evaluate the immune landscape. We also engineered KPL cells with loss of function of EphA2 via CRISPR/Cas9 and tested both single cell clones and multiclonal populations *in vivo*. Unfortunately, we did not observe significant differences in tumor burden between sgEphA2 cell populations and the sgLacz control cells (data not shown). After these observations, we were hesitant to continue this avenue of inquiry, as some of our original data may be confounded by overexpression artifact. Despite this, we do believe a portion of our work reflects true biology, and not all the data are a result of artifact. This sentiment arises from recently published works that highlight EphA2’s role in inhibiting anti-tumor immunity^11^, as well as in monocyte and macrophage function^32–35^, which provide some evidence of veracity of our studies. Discerning which results can be attributed to EphA2 and which to artifact is unfortunately an exceedingly challenging task. Our work not only highlights EphA2’s potential impact on anti-tumor immunity, but also offers a cautionary tale of scientific rigor and reproducibility.

## Acknowledgements

We thank the Vanderbilt Technologies for Advanced Genomics (VANTAGE), Center for Small Animal Imaging, and Flow Cytometry Shared Resource for their assistance.

